# Scene perception-memory pairing extends to superior parietal cortex

**DOI:** 10.64898/2026.05.08.723871

**Authors:** Runhan (Nicole) Tang, Dominika Panek, Leila H. Barkoff, Catriona L. Scrivener, Edward H. Silson, Adam Steel

## Abstract

Visual scene analysis relies on a set of scene-selective regions in posterior cerebral cortex (OPA, PPA, MPA), with a paired memory-responsive counterpart located in anterior (LPMA, VPMA) or overlapping cortex (MPMA). The interaction between these pairs of regions is thought to integrate visual input with mnemonic context. Recently, a fourth scene-perception area in superior parietal cortex (SPA) was identified, with a proposed role in visually-guided navigation. Whether this region also has an anterior paired memory region is currently unknown. Across two independent fMRI datasets (total N=24, 14 females) using static or dynamic stimuli and distinct memory tasks, we show that recalling visual scenes evokes robust responses in a region (referred to here as SPMA) immediately anterior and dorsal to SPA. During resting-state fMRI, SPA preferentially coupled with the other scene-perception areas, while SPMA preferentially coupled with the other place memory areas. At the whole-brain level, seed-based connectivity revealed that SPA sits at the confluence of four processing streams spanning regions implicated in egocentric scene perception, map-based navigation, perspective taking, and goal-directed movement. These findings extend the perception-memory motif associated with visual scene processing to the fourth cortical surface. The widespread anatomical coupling between scene-perception and memory processes reflects the importance of this interaction for flexible, context-grounded navigation.

## Introduction

As we navigate through the world, we rapidly extract information about our visual surroundings to make informed decisions about where we can move and what affordances are available (Bonner and Epstein, 2017; Dwivedi et al., 2020; Bartnik et al., 2025; Mynick et al., 2025; Zamboni et al., 2026). Prior work has linked these abilities to three scene-selective regions on the lateral, ventral, and medial surfaces of posterior cerebral cortex: the occipital place area (OPA)(Hasson et al., 2002; Dilks et al., 2013), the parahippocampal place area (PPA)(Epstein and Kanwisher, 1998), and the medial place area (MPA, also referred to as retrosplenial complex (RSC))(Bar and Aminoff, 2003; Epstein et al., 2007a; Silson et al., 2016b). Each scene perception area is thought to play a unique role(Epstein and Baker, 2019; Dilks et al., 2022), with OPA supporting identification of paths and boundaries (i.e., navigational affordances)(Julian et al., 2016; Kamps et al., 2016; Bonner and Epstein, 2017; Lescroart and Gallant, 2019; Zamboni et al., 2026), PPA supporting scene categorization and landmark recognition(Marchette et al., 2015; Persichetti and Dilks, 2018, 2019; Sun et al., 2021), and MPA supporting map-based navigation(Wolbers and Büchel, 2005; Auger et al., 2012, 2015; Marchette et al., 2014; Persichetti and Dilks, 2019; Steel et al., 2019). Beyond visual analysis, navigation also depends on integrating current visual input with memory of our surroundings. A key question is how the brain connects scene perception with visuospatial memory.

Remarkably, the brain appears to connect perceptual and mnemonic processes by placing brain areas involved in visual scene analysis and visuospatial memory in close anatomical proximity. Each scene perception area is flanked by an anterior, memory-responsive counterpart: the lateral, ventral, and medial place memory areas (LPMA, VPMA, MPMA) (Steel et al., 2021). Using visual imagery to isolate top-down memory processes, recent studies have shown these anterior regions are selectively active during scene recall(Steel et al., 2021; Srokova et al., 2022; Steel et al., 2023, 2024b, 2025), are more functionally connected to the hippocampus than their paired perceptual counterparts, and show unique sensitivity to visuospatial context during both perception and recall(Steel et al., 2021, 2023). The combination of anatomical position and functional connectivity makes the memory-responsive areas ideally positioned to integrate perceptual and mnemonic representations during navigation.

Recently, a fourth scene-perception area in superior parietal cortex, referred to as the posterior intraparietal gyrus scene-selective site (PIGS)(Kennedy et al., 2024) or the superior parietal lobule region (SPL)(Yoon et al., 2025), has been described. Here we use the term superior place area (SPA). Like OPA, SPA responds preferentially to egocentric motion through scenes(Kennedy et al., 2024; Yoon et al., 2025, 2026) and is more strongly connected to OPA than to PPA or MPA at rest(Yoon et al., 2025). Interestingly, this area shows enhanced connectivity to motor regions involved in leg versus hand movements compared with other parietal areas(Yoon et al., 2026). These studies collectively suggest that SPA plays a specialized role in visually-guided navigation.

Does this new scene-selective site in superior parietal cortex follow the same posterior-anterior perception-memory organizational motif as the other scene-selective areas? The answer is not obvious. On the one hand, because SPA is specialized for online visual guidance of ego-centric movement, it may require little access to stored spatial memories. On the other hand, visually-guided navigation is not purely reactive; it involves anticipating paths, planning movements, and integrating current views with prior knowledge of an environment. Because these functions require memory of spatial context, SPA may also have a closely affiliated memory area. In prior work with group analyses, we have observed scene memory related activity in superior parietal cortex in past work (Steel et al., 2021, 2023), and another recent study that adopted this localization approach also reported memory evoked activity in superior parietal cortex (Han and Epstein, 2026). However, these studies did not localize perceptual SPA, and therefore whether superior parietal cortex has a set of paired perception and memory areas remains unknown. Observing the same perception-memory pairing seen on the lateral, medial, and ventral surfaces would suggest that this motif is a general principle of cortex devoted to visual scene analysis, not a feature reserved for regions explicitly involved in map-based navigation.

Here we asked whether the scene perception and memory activity in superior parietal cortex follows a similar topographical arrangement to the lateral, ventral, and medial surfaces. Across two independent fMRI datasets using static (Experiment 1) or dynamic (Experiment 2) stimuli and distinct memory tasks, we find that scene recall evokes a memory-responsive region (SPMA) immediately anterior and dorsal to SPA. Further, SPA sits at the intersection of four cortical streams spanning regions implicated in egocentric scene perception, map-based navigation, perspective taking, and goal-directed movement. Together, these findings extend the posterior-anterior perception-memory motif to a fourth cortical surface and supports a “connect-to-context” framework in which every scene-perception area is tightly coupled to a memory-responsive partner.

## Methods

### Procedure

Here, we investigated the topography and connectivity of scene perception and memory processes in superior parietal cortex in two independent experiments, one collected at the University of Edinburgh and the other at Dartmouth College. We implemented perception and memory tasks in both experiments, which differed in the nature of the stimuli (Figure 1).

**Figure 1.**
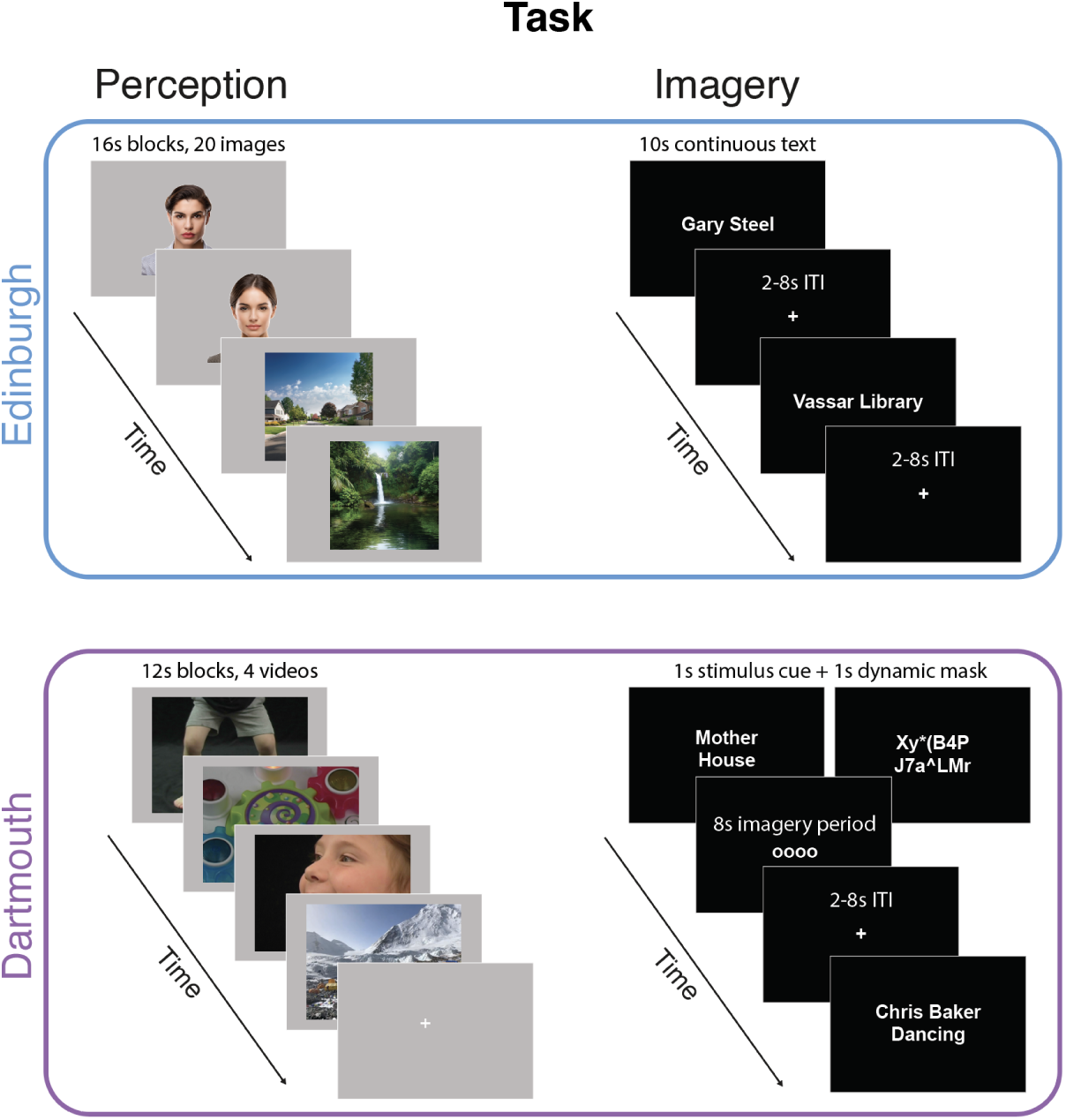
Experimental design. Our study considered two experiments collected at the University of Edinburgh (Experiment 1) and Dartmouth College (Experiment 2). Participants performed perception and memory tasks in both experiments. (upper) In Experiment 1, during the perception task, participants saw a series of unfamiliar face and scene images presented in alternating blocks (16s/block, 20 images; 300ms on, 500ms off); during the imagery task, participants imagined photographs of familiar people or place images. In Experiment 2, during the perception task participants saw a series of 3s videos of unfamiliar bodies, objects, faces, and scenes arranged in blocks (12s/block) with 12s fixation cross intermixed; during the imagery task, participants were cued to imagine 4 different conditions associated with a familiar person: a photograph of the person’s face, the person dancing, having a conversation with the person, or looking around a room in the person’s house. Participants in the Dartmouth experiment also underwent a 10 minute resting-state fMRI scan.

Experiment 1 (Edinburgh) used static images of unfamiliar scenes and faces in the perception task and recall of photographs depicting personally familiar scenes versus people in the memory task. Experiment 2 (Dartmouth) used dynamic scene and face videos in the perception task and recall of familiar scenes versus multiple social stimuli in the memory task. These tasks are described in detail below. To evaluate functional connectivity of the scene perception and memory areas in superior parietal cortex, participants in the Dartmouth experiment also underwent a resting-state fMRI scan.

### Participants

Data collection was approved by each institution’s ethics review boards (Protocol numbers – University of Edinburgh: 420–2122/5 and 420–2122/6; Dartmouth College: 31288). All participants gave informed consent to participate. Prior work showed large effect sizes when evaluating the anterior shift of memory and perception on the cortex’s lateral, ventral, and medial surfaces (Cohens ds > 1.6) (Steel et al., 2021). Based on this effect size (d = 1.6, power = 0.8, alpha = 0.05), 6 participants would be sufficient to detect an effect. Both experiments exceeded this sample size by a factor of two.

#### Experiment 1 (Edinburgh)

This experiment makes use of data reported in(Scrivener et al., 2025), which examined group-average activity of category-selective regions during memory tasks. Here, we consider the spatial location of the place memory areas in posterior cerebral cortex identified in individual participants, including the new area in superior parietal cortex, which had not been previously analyzed. 12 participants (mean age = 27, 3 males) participated in the functional MRI session. All participants were able to perform visual memory based on the Vividness of Visual Imagery Questionnaire (VVIQ = 58.58 mean, range = 38-74). Participants received monetary compensation for their time.

#### Experiment 2 (Dartmouth)

12 participants underwent scanning at Dartmouth College (mean age = 30±8.1 years, 7 males). All participants reported good imagery ability when asked. Measures of trait-level vividness (i.e., VVIQ) were not acquired. Participants received a giftcard as compensation for their time. These data have not been reported previously.

### FMRI tasks

#### Experiment 1 (Edinburgh)

##### Static scene perception localizer

Participants performed two runs of a scene/face localizer task. Participants viewed 20 blocks of 20 unfamiliar scene or face stimuli (images presented 300 ms on, 500 ms off; 16s per block; 10 x 10 degrees of visual angle) while completing a one-back task. Participants indicated a repeated image by making a right-hand button press when the same image was repeated sequentially, which occurred twice per block. There was no fixation/baseline period between blocks.

##### Scene memory task

Prior to scanning, participants provided images of six personally familiar people and six personally familiar places that they could mentally imagine. The names of the individuals/places served as cues for the memory task (visual mental imagery task).

During the imagery task, we instructed participants to create a mental image of the picture linked to the target cue word. This served to standardize their imagery content across stimulus repetitions and encouraged participants to perform imagery rather than recalling general aspects of the people and places. We instructed participants to keep their eyes open and to fixate on the cross during imagery trials.

Participants completed six runs of the mental imagery task, lasting 6 min each. Each imagery cue was presented twice per run (12 total repetitions). One participant only completed four of the six imagery runs.

#### Experiment 2 (Dartmouth)

##### Dynamic scene localizer

Participants performed two runs of a dynamic multi-category visual localizer task. Stimuli were publicly available and used in previous studies(Pitcher et al., 2011) and consisted of 3-second videos of four visual categories: i) children’s faces, ii) bodies parts (hands, feet, torso), iii) toys, and iv) scenes.

Participants saw blocks of stimuli from each condition, with four movies per block (4 videos * 3s/videos = 12s/block, 11.4 x 8.5 degrees visual angle). Participants passively viewed the videos and did not perform any behavioral task. All categories appeared four times per imaging run. We organized runs so that participants saw all categories before any category repeated. Before repeating the category blocks, we included a fixation block (12 s duration) to allow the BOLD signal to return to baseline.

##### Dynamic scene memory localizer

Data from Dartmouth were collected as part of a study investigating spatial and social imagery. Prior to scanning, participants provided the names of six people for whom they could imagine the following four prompts: i) imagine a room in that person’s house, ii) a specific photograph of that person, iii) that person dancing, and iv) having a conversation with that person. These people and prompts served as cues for the spatial versus social imagery task. Here, we only considered the contrast of spatial compared with the mixed social conditions; the results specific to the social imagery conditions will be discussed in a separate report.

In the scanner, we showed participants the names they provided along with one of the following cues: house, photo, dancing, conversation. This indicated which stimulus they should imagine, and the condition they would perform.

For the imagery conditions, we provided the following instructions to participants:

##### House

Imagine a specific room in that person’s house. When you are imagining the room, you should envision that the room has materialized all around you, and you should direct your attention smoothly around the environment as though you were standing in that place.

##### Photo

Imagine a specific photograph of that person. The photograph should only include that person and should focus on their face. Hold that specific photograph in mind as long as the cue appears on screen.

##### Dancing

Imagine that person dancing. You should imagine their body moving dynamically through space in a way that would be natural for them. Focus on their movements and the specific way that they move their body, rather than on their face or expression. If it is impossible for you to imagine them dancing, you could imagine them walking, running, or jumping.

##### Conversation

Imagine having a conversation with that person. You should imagine the specific words that they are saying and the sound of their voice. You should not imagine their face or expression, and instead focus on the sound quality, including their tone and unique vocal inflections.

Simulation trials lasted 10 seconds. Trials began with the name/condition stimulus (1s), followed by a dynamic mask (1s, comprising ascii characters that switched every 200 ms), and the imagery cue (8s, four circles displayed at the center of the screen). We instructed participants to perform mental imagery for the entire duration that the imagery cue was on the screen. Trials were separated by a jittered ITI (4-8s), during which time a fixation cross appeared on the screen. During the ITI, we instructed participants to clear their mind and prepare for the next trial.

Participants underwent 4 runs of fMRI scanning on the social imagery task. Each stimulus x condition pairing was shown once per run (4 stimuli per condition x 4 conditions = 16 trials per run). Each scanning run lasted 280s (140 TRs).

##### Resting-state fMRI

For participants in the Dartmouth dataset, we collected 10 minutes of resting-state fMRI data(Biswal et al., 1995). We chose this quantity based on prior work suggesting pairwise regional functional connectivity in cortex stabilizes between 8-12 minutes(Gonzalez-Castillo et al., 2014; Poldrack et al., 2015). During the resting–state scan, we instructed participants to fixate on a white cross shown over a black background and allow their thoughts to wander, to keep their eyes open, and avoid falling asleep. After scanning, all participants reported compliance with these instructions.

### MRI acquisition

Data from both sites were acquired on Siemens 3T Prisma scanners with a 32-channel head coil. Functional scans were acquired using a multiecho multiband echo planar imaging sequence (TR = 2, TEs = 14.6 ms, 35.79 ms, 56.98 ms, MB factor = 2, acceleration factor = 2, 48 interleaved slices, phase-encoding anterior to posterior, transversal orientation, slice thickness = 2.7 mm, voxel size = 2.7 mm × 2.7 mm, distance factor = 0%, flip angle = 70 degrees) identical to prior studies (Steel et al., 2022, 2023, 2024b; Scrivener et al., 2025; Scrivener and Silson, 2025). A high resolution anatomical image was collected for each participant (Edinburgh: TR = 2.5s, TE = 4.37 ms, flip angle = 7 degrees, FOV = 256 mm × 256 mm × 192 mm, resolution = 1 mm isotropic, acceleration factor = 3; Dartmouth: TR = 2300 ms, TE = 2.32 ms, inversion time = 933 ms, flip angle = 8°, FOV = 256 × 256 mm, slices = 255, voxel size = 1 × 1 × 1 mm). T1-weighted images were segmented, and surfaces were generated using Freesurfer (version 6.0)(Fischl, 2012) and aligned to the fMRI data using align_epi_anat.py and @SUMA_AlignToExperiment(Saad and Reynolds, 2012).

### FMRI Pre-processing

#### Dataset 1 (Edinburgh)

MRI scans were processed using AFNI(Cox, 1996), SPM(Friston et al., 1994), Freesurfer(Fischl, 2012), and SUMA(Saad and Reynolds, 2012). Before pre-processing, the origin of all MRI data was manually reset to the anterior commissure (ac) using SPM display (SPM12). For the structural images, the ac origin was defined in the T1w image and the transformation applied to the T2w image, to ensure that they were aligned to each other. For the EPIs in each run, the origin was defined from the first TR of the second echo of the first run, and applied to all other echoes and scans in the same session. This procedure was to facilitate co-registration of EPIs and structural scans across the two sessions, given that the origins of the acquired data could be far apart in space.

During preprocessing, five dummy scans were first removed from the start of each run (AFNI 3dTcat). Large deviations in signal were removed (3dDespike)(Jo et al., 2013) followed by slice time correction (3dTshift), aligning each slice with a time offset of zero. The skull was removed from the first echo 1 scan (TE = 14.6 ms) and used to create a brain mask (3dSkullStrip and 3dAutomask), as this echo contained the most signal. The first echo 2 TR from the first scan (TE = 35.79 ms) was used as a base for motion correction and registration with the T1 structural scan (3dbucket).

After completing the standard preprocessing, the data were processed using Tedana(Kundu et al., 2012, 2013; DuPre et al., 2019, 2021) to denoise the multi-echo scans (version 0.0.12, using default options). In brief, the optimal combination of the three echoes was calculated, and the echoes were combined to form a single, optimally weighted time series (T2smap.py, included in the tedana package). PCA was applied, and thermal noise was removed using the tedana_orig method. Subsequently, ICA was performed, and the resulting components were classified as signal and noise based on the known properties of the T2* signal decay of the BOLD signal versus noise. Components classified as noise were discarded, and the remaining components were recombined to construct the optimally combined, denoised time series. The Tedana optimally combined and denoised output was then scaled to percent signal change.

#### Dataset 2 (Dartmouth)

Multiecho data processing was implemented based on the multiecho preprocessing pipeline from afni_proc.py in AFNI(Reynolds et al., 2024). Signal outliers in the data were attenuated (3dDespike)(Jo et al., 2013). After despiking, motion correction was calculated based on the second echo, and these alignment parameters were applied to all runs. We then optimally-combined and denoised the data using multiecho ICA (ME-ICA) denoising (tedana.py, version 24.0.0)(Kundu et al., 2012, 2013; DuPre et al., 2019, 2021) described above, with the Kundu decision tree for PCA/ICA component classification.

For task fMRI data, following tedana denoising, we applied a 3mm gaussian kernel (3dBlurInMask) in the volume. The time series were then normalized to percent signal change and submitted to 3dDeconvolve (see below). Deconvolved data were then mapped to each participants’ surface (3dVol2Surf)

For resting-state fMRI data, we applied additional preprocessing steps to the data after tedana denoising. Specifically, we used 3dTProject to remove motion, global signal (GSR), and apply a bandpass filter (0.008-0.11 hz). To calculate the global signal, we extracted the average activity from a whole brain mask (grey matter, white matter, and CSF). We applied global signal regression based on evidence showing its effectiveness in removing residual effects of motion and physiological noise(Power et al., 2012, 2017, 2018). Although GSR can induce spurious negative correlations(Saad et al., 2012), our primary interest was differentiating patterns of positive connectivity. We therefore prioritized the beneficial effects of global signal regression for denoising because of our limited data quantity.

Following denoising, we mapped the resting-state data to the individual participant surfaces (3dVol2Surf) and smoothed them with a 3mm gaussian kernel (SurfSmooth).

### FMRI data analysis

#### Task deconvolution and region of interest definition - Edinburgh

The deconvolution procedure for these data are described in the earlier publication(Scrivener et al., 2025). Below we briefly describe the procedure for the reader’s convenience.

##### Static scene perception

To localize the scene perception areas (OPA, PPA, MPA, and SPA) we implemented a generalized linear model (3dREMLfit)(Olszowy et al., 2019) for the scene/face localizer scans. We modeled each category’s activation using a block design convolved with a 16-s GAM basis function (GAM: 8.6, 0.547, 16, 3dDeconvolve). The final generalized linear model included these two task regressors. We included 6 parameters of motion as regressors of no interest. During regression, we estimated the temporal autocorrelation of each voxel’s time series using residual maximum likelihood (REML) estimation to find the best-fit ARMA(1,1) model for the time series correlation matrix. We projected the resulting maps to each individual’s cortical surface (3dVol2Surf) and smoothed them with a 2mm gaussian kernel.

Scene perception areas (OPA, PPA, MPA, and SPA) were defined for each participant by contrasting activity from scene versus face blocks (t-stat > 2.79 (p<0.005), surface cluster size (k) > 20). SPA was defined as the peak in selectivity near the dorsal extent of the parieto-occipital sulcus and posterior intraparietal gyrus (as described in prior work(Kennedy et al., 2024; Yoon et al., 2025)).

##### Static scene memory

To localize the place memory areas (LPMA, VPMA, MPMA, and SPMA), we implemented a generalized linear model (3dREMLfit)(Olszowy et al., 2019) with each imagery category’s activation modeled using a BLOCK basis function (BLOCK(9.5,1), 3dDeconvolve) aligned to the onset of the imagery period. The final generalized linear model included these two task regressors. We included 6 parameters of motion as regressors of no interest. During regression, we estimated the temporal autocorrelation of each voxel’s time series using REML estimation to find the best-fit ARMA(1,1) model for the time series correlation matrix. We projected the resulting maps to each individual’s surface (3dVol2Surf) and smoothed them with a 2mm gaussian kernel.

We defined the place memory areas for each participant by contrasting activity from place versus people imagery trials (t-stat > 2.79 (p<0.005), surface cluster size > 20). In many subjects, this imagery contrast revealed a large cluster in superior parietal cortex with multiple peaks of selectivity. We considered the most posterior of these peaks to be the SPMA region, and we defined SPMA by separating the cluster into its distinct peaks based on the topography of the t-statistic. These subdivisions were made by two authors independently (N.T. and D.P.), and any disparity in the definition was resolved by a third author (A.S.). Importantly, because we considered the posterior cluster SPMA, subdividing this larger memory cluster into its subdivisions would make identifying a distinction between SPA and SPMA less likely.

#### Task deconvolution and region of interest definition - Dartmouth

##### Dynamic scene perception

To localize the scene perception areas (OPA, PPA, MPA, and SPA) we implemented a generalized linear model (3dREMLfit)(Olszowy et al., 2019) for the multicategory localizer scans. We modeled each category’s (scene, face, object, body) activation using a block design convolved with a 12-s GAM basis function (GAM: 8.6, 0.547, 12, 3dDeconvolve). The final generalized linear model included these four task regressors. We included 6 parameters of motion as regressors of no interest. During regression, we estimated the temporal autocorrelation of each voxel’s time series using REML estimation to find the best-fit ARMA(1,1) model for the time series correlation matrix. We projected the resulting maps to each individual’s surface (3dVol2Surf). Because we applied smoothing to these data before deconvolution, no further smoothing was applied.

Scene perception areas (OPA, PPA, MPA, and SPA) were defined for each participant by contrasting activity from scene versus face blocks (t-stat > 2.79 (p<0.005), surface cluster size > 20). SPA was defined as the peak in selectivity near the dorsal extent of the parieto-occipital sulcus and posterior intraparietal gyrus(Kennedy et al., 2024; Yoon et al., 2025).

##### Dynamic scene memory

To localize the place memory areas (LPMA, VPMA, MPMA, and SPMA), we implemented a generalized linear model (3dREMLfit)(Olszowy et al., 2019) with a regressor for each simulation condition (house, photograph, conversing, dancing) modeled with a GAM basis function (GAM: 8.6, 0.547, 8, 3dDeconvolve) aligned to the onset of the imagery period. The final generalized linear model included these four task regressors. We included 6 parameters of motion as regressors of no interest. During regression, we estimated the temporal autocorrelation of each voxel’s time series using REML estimation to find the best-fit ARMA(1,1) model for the time series correlation matrix. We projected the resulting maps to each individual’s surface (3dVol2Surf). Because we applied smoothing to these data before deconvolution, no further smoothing was applied.

We defined the place memory areas for each participant by contrasting activity from place versus all other category simulation trials (t-stat > 2.79 (p<0.005), surface cluster size > 20). In many subjects, this imagery contrast revealed a large cluster in superior parietal cortex with multiple peaks in selectivity. We considered the most posterior of these peaks to be the SPMA region, and we defined SPMA by separating the cluster into its distinct peaks based on the topography of the t-statistic. These subdivisions were made by two authors independently (N.T. and D.P.), and any disparity in the definition resolved by a third author (A.S.). Importantly, because we considered the posterior cluster SPMA, subdividing this larger memory cluster into its subdivisions would make identifying a distinction between SPA and SPMA less likely.

#### Topographic shift analysis

For both datasets, we analyzed the topographic shift between the scene perception and memory clusters using the approach adapted from prior work(Steel et al., 2021). After defining individual regions of interest, we quantified the anterior displacement of perception and memory areas by calculating the center of mass for the paired scene perception/place memory selective area on each surface. We then compared the distance between the center of mass for perception and memory in the y-dimension (posterior-anterior) using a linear mixed effects model with Task (Perception, Memory) and Hemisphere (LH/RH) as factors and participant as a random effect for each surface separately. We used linear mixed effects modeling to account for participants in whom we could only identify regions in one hemisphere more effectively. Because there was no significant interaction between Task and Hemisphere (all ps>0.10), we present the data collapsed across hemispheres.

We calculated the angular consistency of the shift for each region using a Rayleigh’s test. We transformed the coordinates from cartesian to polar; for this analysis we only considered the principle orientation of the dorsal surface (Y and Z).

To examine whether the proportion of shared vertices differed across surfaces, we compared the overlap between the category-selective perception and memory areas on each surface. We calculated the Jaccard Index (proportion of shared vertices between the perception and memory areas as a function of total vertices), and compared them using a linear mixed effects model with Surface (Medial, Ventral, Lateral, Dorsal), Hemisphere (LH/RH) as fixed effects, and participant as a random effect.

#### Exploratory group analysis

To quantify the overall pattern of brain activity evoked by perception and memory of scenes, we performed group analyses for the experiments collected at Edinburgh and Dartmouth. We conducted four separate 1-sample t-tests for the relevant contrast maps for perception (Edinburgh: Scene v Face images; Dartmouth: Scene v Face videos) and memory (Edinburgh: Place v People imagery; Dartmouth: House v mean across social conditions) tasks from both experiments. We then qualitatively compared the location of the significant areas (t > 3.3, p < 0.008, k > 40) between the perception and imagery tasks in each dataset. Based on the importance of retinotopic coding to visual scene analysis, to aid interpretation we compared the group activation maps to the known retinotopic maps in superior parietal cortex using the Wang atlas(Wang et al., 2015).

#### Resting-state functional connectivity analyses

To investigate how the SPA fits into the broader organization of cerebral cortex, we performed two different functional connectivity analyses. To ensure that all regions were equally represented and not biased by region size, we constrained all regions of interest to the 300 vertices closest to the center of mass of the original area defined above. All original regions had greater than 300 vertices.

##### Connectivity among the scene perception and place memory areas

To examine how the superior parietal scene perception and memory areas (SPA v SPMA) was functionally related to the originally described scene perception and memory areas, we compared the pairwise correlation of resting-state activity among all regions (perception areas: OPA, PPA, MPA, SPA; memory areas: LPMA, VPMA, MPMA, SPMA). We extracted the resting-state fMRI time series from the 300 vertex constrained center of mass regions of interest. Next, to isolate the unique variance associated with each pairwise connection (Smith et al., 2013), we performed a partial correlation among all region pairs (partialling out the variance associated with other regions). We then compared the affiliation between SPA and SPMA with the perception and memory areas using a linear mixed effects model with Hemisphere (LH/RH), Superior parietal region (SPA, SPMA), Surface (Lateral, Medial, Ventral), and the task used to define the regions (Task: Perception, Memory) as factors and participant as a random effect. We observed no effect or interaction with the hemisphere, so data are presented averaged across hemispheres.

##### Connectivity to other cortical areas

As an exploratory analysis we used whole-brain seed-based connectivity to investigate how SPA connects with the rest of the brain. We extracted resting-state fMRI time series from the constrained SPA and SPMA ROIs and calculated, separately for each seed and hemisphere, the correlation between the seed time series and the time series at every cortical vertex. To aggregate data across participants, we created binary masks of the top 10% most connected vertices to the SPA perception and memory regions. This method of percentile based thresholding has been used by prior resting-state studies to derive individualized connectivity maps(Dickie et al., 2018). We then averaged these binary masks together to derive a whole-brain probability map of SPA connections for all participants. We conducted this analysis for each hemisphere separately.

## Results

### The posterior-anterior perception-memory motif extends to superior parietal cortex

We began by asking whether scene perception and memory (visual imagery) tasks activate dissociable regions within superior parietal cortex. In Experiment 1, we considered perception and visual memory tasks that closely matched prior work: i) perception of static scene images compared with face images (Methods: Static scene perception) and ii) visual imagery of familiar places compared with visual imagery of familiar people (Methods: Static scene memory). In Experiment 2, we considered multiple social demands during both perception and recall trials: i) perception of videos of panning scenes versus videos of children’s faces (Dynamic scene perception), and ii) visual recall of familiar places compared with dynamic social simulation (Dynamic scene memory, see Methods).

For each participant, we defined regions of interest for SPA and SPMA in each hemisphere. We also defined regions of interest for the known scene perception areas and place memory areas on the medial, ventral, and lateral surfaces. Consistent with previous work, we quantified the shift in the center of mass from scene perception to place memory in each participant (in millimeters on the cortical surface), consistency of the shift direction, and the overlap between the scene perception and memory areas(Steel et al., 2021, 2025). For both scene perception and memory recall ROIs were drawn using a vertex-wise threshold of t > 2.79 (p<0.005) and a cluster size of 20 connected surface vertices. Notably, we considered SPMA contingent on observing SPA in a participant.

We observed a consistent topographic shift of place memory compared with scene perception in superior parietal cortex. Observing individual participants’ activation revealed a striking pattern: SPA formed the posterior border of a large cluster of scene memory activation (four example participants from Experiment 2 are depicted in Figure 2A). Across participants, this large cluster of memory activity had up to 4 distinct peaks anterior and dorsal to SPA, spanning superior parietal lobule and the marginal ramus of the cingulate.

**Figure 2.**
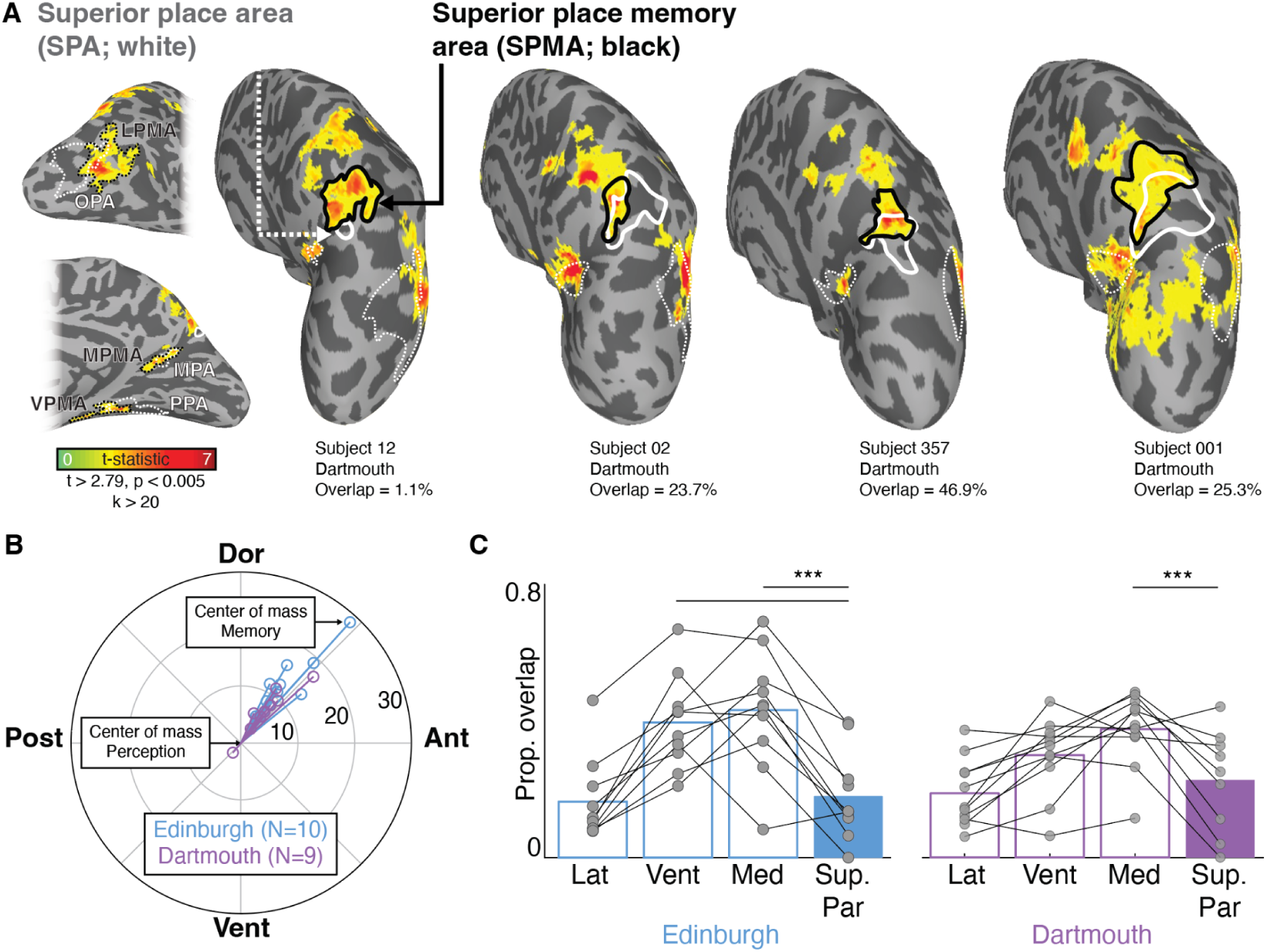
Anterior shift of scene memory versus perception activity in superior parietal cortex. A. Example scene perception (white outline) and memory area (black outline) in superior parietal cortex in four participants. Memory activity for scene versus social imagery conditions is shown in hot colors. Lateral and medial surfaces are shown for subject 12 showing typical locations of the scene perception (white dotted lines) and place memory areas (black dotted lines). The display threshold (t>2.79) was used to define the regions of interest for subsequent analyses. B. Consistent anterior and dorsal shift of place memory area versus perception area in superior parietal cortex. Perception and memory areas were drawn in individual participants in both experiments based on a vertexwise t>2.79 (p<0.005). Center of the polar plot depicts the center of mass of perceptual area, lines depict the shift in each participant in millimeters. C. Superior parietal perception and memory clusters have limited overlap compared with medial and ventral surfaces. Data points show the Jaccard Index (proportion of overlapping vertices as a function of total vertices) for perception and memory areas on each surface for each participant. Data in B and C are averaged across hemisphere.

Because the number and distribution of local maxima within the superior parietal cortex during the memory task were idiosyncratic across participants, we defined SPMA by separating the cluster into its distinct peaks based on the topography of the t-statistic maps (tracing local minima between cluster peaks). As a conservative definition of SPA paired region, we considered the most posterior peak to be the SPMA. That is, we selected the peak most likely to minimise the anterior shift. Two authors independently (N.T. and D.P.) made these definitions, and any disparity in their definition of SPMA was resolved by a third author (A.S.). Importantly, because we considered the most posterior cluster SPMA, subdividing this larger memory cluster into its subdivisions would make identifying a distinction between SPA and SPMA less likely.

Many participants also had a peak of scene memory activity at the inflection point of the marginal ramus of the cingulate, consistent with the location of a region called “Places-2” in our prior work(Silson et al., 2019b; Scrivener et al., 2025; Scrivener and Silson, 2025). We considered this to be an independent area, and therefore we did not label it as SPMA in any participant. Because this paper focuses on the topographic adjacency of perception and memory-related activity, defining this isolated memory cluster in individuals falls outside the scope. Future studies defining this marginal ramus scene memory cluster within individuals could provide further insight into the role of superior parietal cortex in memory-related processes.

Using this approach, we successfully localized SPA and SPMA bilaterally in most participants. In Experiment 1, we localized SPA in 10/12 participants; all participants with SPA had an associated memory area (10/10). In Experiment 2, we localized SPA in 11/12 participants; among these 11 participants, we observed SPMA in 9/11 participants bilaterally. Note that the smaller number of participants with SPMA in Experiment 2 could be due to the complexity of the memory task and the smaller number of scene memory trials in this task. This suggests a robust response during memory within superior parietal cortex, complementing the description of SPA in perceptual tasks(Kennedy et al., 2024; Yoon et al., 2025).

Across both studies, all but one participant showed an anterior shift of perception versus memory activation in superior parietal cortex (Figure 2B; Experiment 1: 10/10; Experiment 2: 8/9; across experiments = 94.73%). At the group-level, this anterior shift was significant in both experiments, with a mean anterior-posterior shift of 7.73 (range: 0.75-23.1) and 4.48 (−1.45-12.71) millimeters in Experiment 1 and 2, respectively, (Linear mixed effects model, Main effect of task – Experiment 1: F(1,36)=11.58, p=0.0016, eta^2^=0.243; t(9)=4.86, p=0.00090, d=1.378; Experiment 2: F(1,36)=10.148, p=0.0029, eta^2^=0.215; t(8)=3.41, p=0.009, d=1.136). There was no interaction between hemisphere and task in either experiment (LME, Hemisphere x Task – Experiment 1: F(1,36)=0.009, p=0.92, eta^2^<0.01; Experiment 2: F(1,36)=0.016, p=0.89, eta^2^<0.01). This shift was consistently anterior and dorsal, which we confirmed based on a Rayleigh’s test of uniformity (Experiment 1: z=7.934, p<0.0001; Experiment 2: z=6.28, p=0.00061).

In addition to the superior parietal cortex, we confirmed that these new samples of participants showed the anterior shift of memory versus perception across the other cortical surfaces (Steel et al., 2021, 2025). All regions of interest were present in at least one hemisphere in 10/12 participants in Experiment 1 and 11/12 participants in Experiment 2. Consistent with prior reports, we observed the anterior shift of memory compared to perception activity for scenes in both Experiments on the lateral and ventral surfaces (Linear mixed effects model, Main effect of task and follow up t-tests: Experiment 1 – Lateral: F(1,36)=130.63, p<0.0001, eta^2^=0.789, t(9)=11.75, p<0.0001, d=3.72; Ventral: F(1,36)=33.55, p<0.0001, eta^2^=0.482, t(9)=7.11, p<0.0001, d=2.25; Experiment 2 – Lateral: F(1,41)=87.85, p<0.0001, eta^2^=0.682, t(10)=12.13, p<0.0001, d=3.66; Ventral: F(1,41)=16.67, p=0.0002, eta^2^=0.289, t(10)=4.17, p=0.0019, d=1.26). On the medial surface, we observed no anterior shift on the medial surface in Experiment 1, and a tendency for a posterior shift in Experiment 2 (Experiment 1: F(1,36)=1.26, p=0.27, eta^2^=0.034; t(9)=0.75, p=0.47, d=0.23; Experiment 2: F(1,42)=6.53, p=0.01, eta^2^=0.135, t(10)=-1.08, p=0.304, d=0.33). This further strengthens the evidence for an anterior shift on the lateral and ventral surfaces (Steel et al., 2025).

All of the place memory areas tend to be anteriorly shifted compared to their perceptual counterparts. However, the degree of overlap between these areas differs by cortical surface, with the lateral, ventral, and medial surfaces ordered least to greatest(Steel et al., 2021). How would SPA and paired memory area fit within this range? Intriguingly, the proportion of overlap between perception and memory activation in superior parietal cortex was similar to the OPA and LPMA on the lateral surface (Figure 2C). When we compared the Jaccard Index (proportion of overlapping vertices vs total) across all surfaces (medial, ventral, lateral, superior parietal), we found the proportion of overlapping vertices was significantly lower in the superior parietal versus the medial and ventral surfaces in Experiment 1 (LME, Main effect of Surface – Experiment 1: F(3,73)=13.144, p<0.0001, eta^2^=0.354; Follow-up t-tests – Experiment 1: Sup. Par. vs Ventral t(9) = 3.58, p = 0.0059, d = 1.13, Sup. Par. vs Medial t(9) = 5.72, p = 0.00029, d = 1.80, Sup. Par. vs Lateral t(9) = −0.32, p = 0.75, d = −0.1018, Lateral vs Ventral t(9) = −6.85, p < 0.0001, d = 2.17, Lateral vs Medial t(9) = −5.69, p = 0.00030, d = 1.7989, Ventral vs Medial t(9) = −0.63, p = 0.55, d = −0.199). In Experiment 2, we found significantly lower overlap between the place memory and scene perception areas in superior parietal cortex compared with the medial surface, and a numerically lower overlap compared with the ventral surface (although this was not statistically significant) (LME, Main effect of Surface – Experiment 2: F(3,73)=4.467, p=0.006, eta^2^=0.153; Follow-up t-tests – Sup. Par. vs Ventral: t(8) = 1.902, p = 0.094, d = 0.634, Sup. Par. vs Medial: t(8) = 3.88, p = 0.005, d = 1.293, Sup. Par. vs Lateral: t(8) = −0.603, p = 0.563, Lateral vs Ventral, t(10) = −3.56, p = 0.005, d = 1.073, Lateral vs Medial: t(10) = −4.478, p = 0.001, d = 1.350, Ventral vs Medial: t(10) = −2.310, p = 0.044, d = 0.696).

### Scene perception and memory activity border parietal retinotopic maps

Analysis of individual participants’ activity provided strong support for the distinction between the perception and memory areas in superior parietal cortex. To highlight the consistency across experiments and situate these findings with prior literature, we performed an exploratory group analysis of the scene perception and memory tasks separately for both experiments (Figure 3). All participants (including those in whom we could not localize SPA or SPMA individually) were used in this group analysis. We compared the group activation maps to the known retinotopic maps from the Wang atlas (Wang et al., 2015) based on the importance of retinotopic coding in perception and memory (Silson et al., 2019b; Breedlove et al., 2020; Groen et al., 2021; Favila et al., 2022; Angeli et al., 2024; Steel et al., 2024b, 2024a; Scrivener and Silson, 2025).

**Figure 3.**
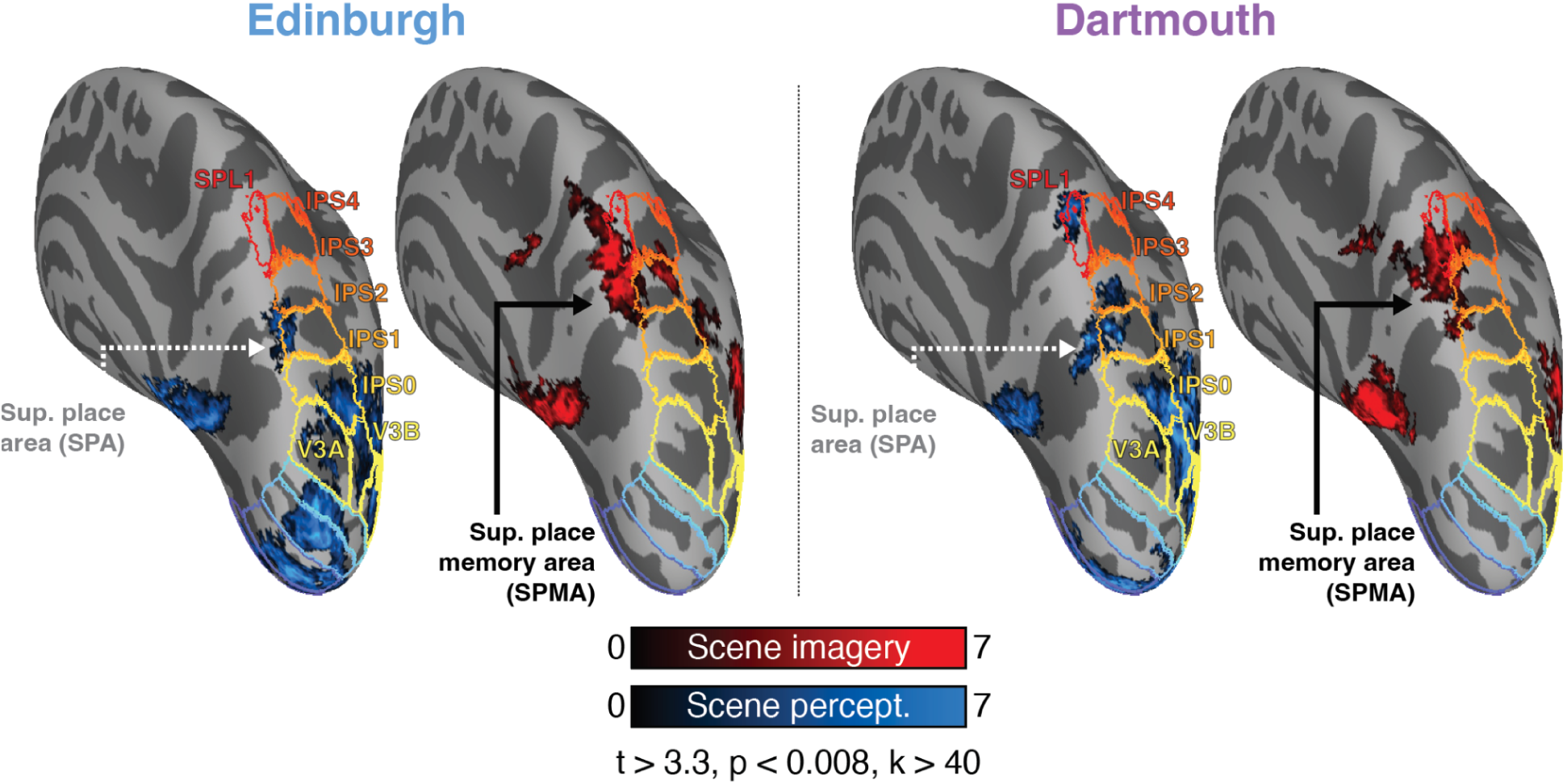
Scene perception and place memory activity occur in distinct locations in superior parietal cortex. We conducted a group analysis of the scene perception (blue) and memory (red) conditions for each experiment and compared these results to probable locations of known retinotopic maps in intraparietal sulcus (defined using (Wang et al., 2015)) (group analysis thresholded at t>3.3, p<0.008, k>40). In both experiments, scene perception activity occurred within the retinotopic map IPS1, while scene memory activity straddled the medial boundary of maps IPS2, IPS3, and SPL1. In both experiments, we observed additional scene memory activity at the inflection point of the marginal ramus of the cingulate, consistent with the region “Places-2”(Silson et al., 2019b; Scrivener et al., 2025; Scrivener and Silson, 2025).

Our group analysis revealed highly consistent areas of activation across experiments, with scene memory activity anteriorly shifted relative to perception. For scene-selective perceptual activity, we observed a cluster in the medial portion of IPS1 at the dorsal extent of the parieto-occipital sulcus, consistent with the location of SPA(Kennedy et al., 2024; Yoon et al., 2025). At our threshold (t > 3.3, p < 0.008, cluster size > 40), this cluster minimally overlapped with IPS2. OPA overlapped with V3B, LO1, LO2, and IPS0 (Silson et al., 2016a; Scrivener et al., 2024), and was separated from SPA by non-significant vertices in the lateral portion IPS1. In Experiment 2, in addition to the group SPA cluster, we observed another patch of scene-selective perceptual activity in superior parietal cortex within the anterior portion of the retinotopic map SPL1. Thus, despite Experiments 1 and 2 using static versus dynamic stimuli, we observed common patterns of scene-selective activity in superior parietal cortex, suggesting robust localization of SPA regardless of stimulus type.

Like the scene perception activation, scene memory activity was consistent across experiments and linked to IPS retinotopic maps. We observed a cluster of scene memory activity anterior to IPS1, spanning the medial portion of IPS2, IPS3, and the posterior portion of SPL1. Importantly, this group SPMA memory cluster straddled the medial edge of these maps, partially overlapping with these maps and extending further medially into precuneus on the medial wall. Notably, at our threshold (t > 3.3), in Experiment 2 the group SPMA terminated posterior to the second scene perception cluster within anterior SPL1, indicating a possible distinction in perception and memory activity within the SPL1 retinotopic map.

In addition to this SPMA cluster, we observed another patch of memory-related activity anterior and medial to SPMA. This cluster, likely corresponding to the Places-2 memory cluster(Silson et al., 2019b; Scrivener and Silson, 2025; Scrivener et al., 2025) or anterior precuneus(Zhang et al., 2025), fell at the inflection point of the marginal ramus of the cingulate on the medial wall and did not overlap with any retinotopic maps. Thus, medial parietal cortex appears to be tiled by three place memory selective regions: a ventral region within the parieto-occipital sulcus (MPMA/Places-1), a dorsal region located within the superior parietal lobule (SPMA), and an anterior region at the inflection point within the cingulate ramus.

### Place memory areas maintain selectivity during visual perception

Our prior analysis shows that recalling personally-familiar places activates brain areas anterior to the scene perception areas. A related question is whether the place memory areas retain their selectivity when perceiving visual scenes. To address this question, we extracted the activity estimates (beta-values) from the place memory areas (including SPMA in the dynamic localizer task collected during Experiment 2 (Dartmouth). We focused on the Dartmouth dataset because this experiment included rest blocks (12s), allowing us to assess the activation during perception versus a fixation baseline. To equate SNR and signal homogeneity across the regions, we constrained these regions to the 300 vertices closest to each region’s center of mass.

We found that the place memory areas activated in a relatively selective manner during scene perception, consistent with the partial overlap between the scene perception and place memory areas (Figure 4). Across all ROIs, we observed a Main effect of Category (LPMA: F(3,76)=12.96, p<0.001, eta^2^=0.334; VPMA: F(3,76)=33.973, p<0.001, eta^2^=0.573; MPMA: F(3,76)=20.708, p<0.001, eta^2^=0.437; SPMA: F(3,68)=8.931, p<0.001, eta^2^=0.283). Critically, viewing scenes evoked greater activity compared with all other categories in LPMA, VPMA, and MPMA (ts>3.41, ps<0.04). The notable exception was SPMA, where we observed a significant difference between activity during scene perception compared with faces (t(9)=5.395, p=0.003, d=1.70) and bodies (t(9)=3.986, p=0.019, d=1.26), but not objects (t(9)=1.934, p=0.511, d=0.612). This suggests that although these regions are preferentially active during memory tasks, they retain their selectivity when perceiving dynamic visual scenes.

**Figure 4.**
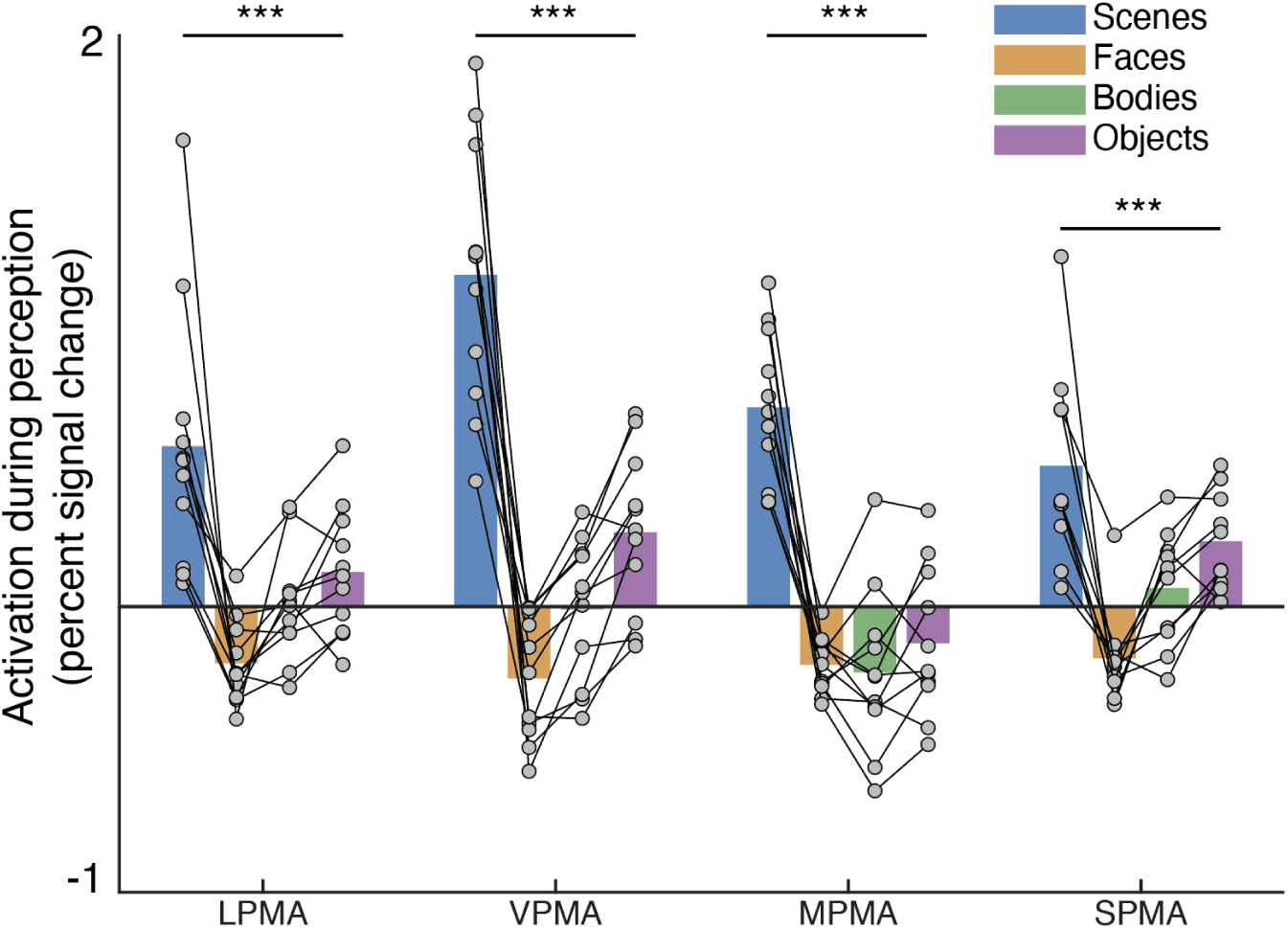
The place memory areas activate in a relatively selective manner to visually-presented scenes. We extracted activation (beta-values) during the dynamic perception task from the place memory areas in participants from Study 2 (Dartmouth). All place memory areas showed greater activity during scene perception compared with other categories (faces and bodies), although SPMA was not significantly different from objects. This is consistent with the general preference for spatial information, even in the absence of memory recall demands.

### Connectivity dissociates superior parietal scene perception and memory areas

Analyzing co-activation of brain areas can provide insight into their function, because co-activation patterns at rest often reflect contributions to similar cognitive/functional processes(Biswal et al., 1995). When watching a video meant to convey naturalistic scene understanding, the scene perception and memory areas on the lateral, ventral, and medial surfaces form distinct functional networks(Steel et al., 2021). Here, we asked whether this distinction between the scene perception and memory areas was also true during rest, and whether the scene perception and memory areas in superior parietal cortex would preferentially associate with regions defined in the same task.

We hypothesized that activity of SPA would be more correlated with the scene perception areas (OPA, PPA, and MPA), while SPMA would be more correlated with the memory areas (LPMA, VPMA, and MPMA). We tested this hypothesis in resting-state fMRI data collected in the Experiment 2 (Dartmouth) participants in whom we could define all perception and memory areas (N=9). This sample size was sufficient to detect a large effect (Cohen’s d > 1.2, 80% power, alpha = 0.05), which is consistent with the functional connectivity between scene perception and place memory areas detected in our prior study (Cohen’s ds > 2.39)(Steel et al., 2021). As above, to equate SNR and signal homogeneity across the regions, we constrained these regions to the 300 vertices closest to each region’s center of mass. We extracted the resting-state fMRI time series from these constrained ROIs, and calculated the partial correlation among all region pairs (while partialling out variance shared with all of the regions). After constraining to the 300 vertices around the centers of mass, ROIs minimally overlapped (all ROIs < 50% overlap) and results did not differ if only unique vertices were included.

Two principles appeared to organize the functional relationship among the perception and memory areas. The first principle was anatomical pairing: we observed high connectivity strength between anatomically paired perception and memory areas (Figure 5A, secondary-diagonal). The second principle was perceptual vs mnemonic region: aside from the paired region, all other across network connections (i.e., perception to memory area pairs) was lower than within network connections. Thus, these regions’ resting-state activity captures their association during the localizer tasks.

**Figure 5.**
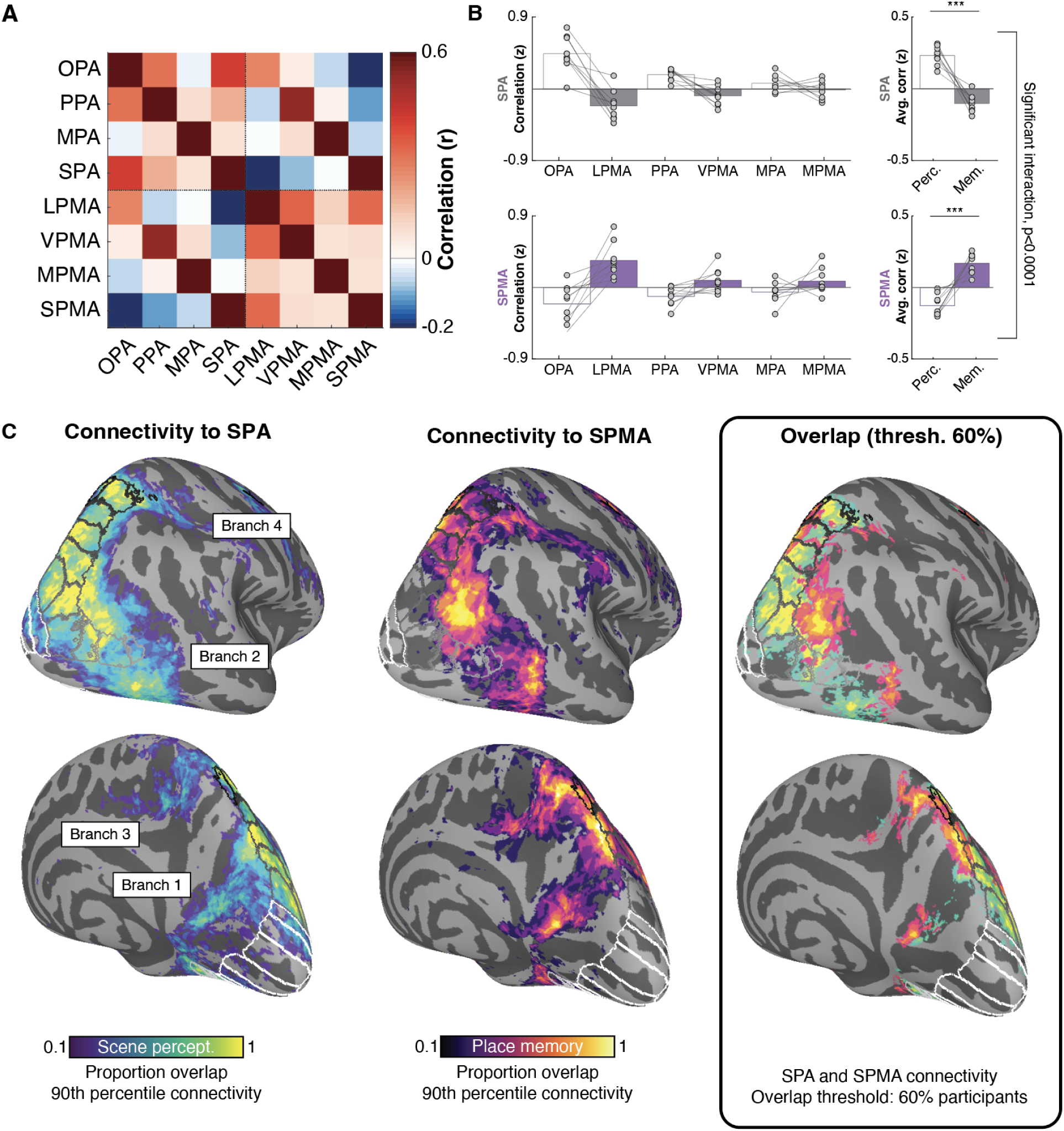
Functional connectivity during rest dissociates perception and memory areas in superior parietal cortex. A. Pairwise partial correlation(Smith et al., 2013) between scene perception areas (OPA, PPA, MPA, SPA) and place memory areas (LMPA, VPMA, MPMA, SPMA), providing an estimate of the unique variance explained at each connection. B. Differential connectivity between SPA and SPMA with the other scene perception and place memory areas. SPA was more connected with other scene perception areas, while SPMA is more connected with the place memory areas. C. Parallel branching connectivity streams within the superior parietal cortex emanating from SPA (left) and SPMA (middle). In each participant, we calculated whole-brain connectivity of SPA and SPMA, then created binarized masks of the 90th percentile of connected vertices. We aggregated these masks across participants to create a probabilistic map of SPA and SPMA connectivity. This analysis revealed parallel streams of connectivity spanning superior parietal cortex and intraparietal suclus. Connectivity from SPA largely follows IPS, with branches emanating 1) posterior medially (toward MPA/MPMA), 2) posterior ventrally (toward OPA/LPMA), 3) dorsal medially (toward marginal ramus of the cingulate, places-2), and 4) dorsal laterally (toward parietal area PF). Connectivity from SPMA borders these connectivity streams on the lateral and medial sides, with peaks of connectivity falling anterior to the peaks of SPA connectivity at each branch’s terminus. The right panel depicts the SPA and SPMA probability maps, showing vertices that exceeded a 60% overlap across participants.

With respect to perception and memory areas in superior parietal cortex, SPA and SPMA were affiliated with the other scene perception and memory areas at rest. We observed a double dissociation in the connectivity between the superior parietal perception and memory areas with the other surfaces’ memory and scene perception areas (Figure 5B; LME – Superior parietal area x SPA/PMA interaction: F(1,96)=135.7, p<0.001, eta^2^=0.586). This was driven by preferential connectivity of SPA to the other perceptual versus memory areas, and vice versa for SPMA (Difference SPA v SPMA connectivity for perception versus memory areas: t(8)=4.165, p=0.003, d=1.388; Perception v memory area connectivity – SPA: t(8)=8.591, p<0.001, d=2.864; SPMA/PIGS: t(8)=-6.715, p<0.001, d=2.238). Notably, all participants showed this double dissociation. This is consistent with the hypothesis that the scene perception and place memory areas constitute dissociable functional networks, including the new regions within superior parietal cortex.

### Distinct medial and lateral connectivity streams originate from the superior parietal perception area

Analysis of the topography of scene perception and memory activation in superior parietal cortex showed that SPA formed the posterior boundary of a larger swath of memory-evoked activity within this region. This led us to ask how SPA and SPMA functionally connect with the rest of the cortex, particularly superior parietal cortex anterior to SPA.

To address this question, we conducted an exploratory seed-based connectivity analysis of SPA and SPMA. We extracted resting-state fMRI time series from the individually-defined SPA (n=10) and SPMA regions (n=9) and calculated the whole brain correlation map with SPA (separately for each hemisphere). We then created a binary mask of the top 10% most connected vertices for each participant and aggregated these binary masks across participants to create a vertex-wise probability map. Although we did not explicitly stipulate a positivity constraint, only positive connectivity values were included in the 10% connectivity mask. Using this binary mask approach (versus other group analysis approaches like t-testing) prevents bias from outlying connectivity values within individual participants and ensures that all participants contribute equally to the group-level map.

This analysis revealed that connectivity from SPA and SPMA form parallel connectivity pathways extending throughout parietal cortex. SPA sits at the posterior confluence of four anatomically distinct branches of preferential connectivity extending into the dorsal parietal cortex. These branches are bordered on their medial and lateral sides by areas with high connectivity with SPMA. Figure 5C shows the proportion of overlapping participants connectivity maps from SPA (Left), SPMA (middle), and the overlap between these maps thresholded at 60% overlapping participants (right).

Ventrally, connectivity from SPA follows two distinct branches. Branch 1 runs laterally, connecting SPA to OPA. Branch 2 runs medially, connecting SPA to MPA/MPMA via the parieto-occipital sulcus. These connectivity streams converge at SPA and form a single stream moving anterior and dorsally, through SPMA toward superior parietal cortex. Both branches are bordered by increased connectivity with SPMA, the lateral stream beginning in LPMA, and the medial stream beginning in MPA/MPMA.

Dorsally, the connectivity stream emanating from SPA splits into two additional dorsal medial and lateral branches. In the group average map, the bifurcation appeared within the superior aspect of the marginal ramus of the cingulate and the retinotopic map SPL1. At the bifurcation point, Branch 3 dove down the medial wall toward the inflection point of the marginal ramus of the cingulate. Branch 4 extended ventrally and laterally into PF complex and area PFt in lateral parietal cortex(Glasser et al., 2016). The end points of branches 1-3 have recognized roles in navigation-related processes (Marchette et al., 2014; Persichetti and Dilks, 2018; Zhang et al., 2025), and area PF/PFt has recently been identified as a region of a larger brain network related to navigation(Zhang et al., 2025). Connectivity from SPMA followed these branches toward their termini, paralleling their medial and lateral trajectories. This positions the superior parietal cortex as an anatomical hub connecting disparate cortical areas to facilitate visually-guided navigation.

In addition to these parietal connectivity pathways in superior parietal cortex, we observed three other loci of increased SPA and SPMA connectivity distributed in cortex. Interestingly, these areas tended to be at the anterior edges of retinotopic maps(Wang et al., 2015), and at each location, SPMA connectivity fell at the anterior edge of the SPA connectivity cluster. One patch of connectivity in medial ventral temporal cortex, encompassing the retinotopic maps TE1&2 and PHC1&2, consistent with the location of PPA and VPMA. A second patch of connectivity was in lateral occipitotemporal cortex, anterior to the retinotopic maps MT/hMT. Finally, we found a third patch in lateral prefrontal cortex, anterior to the frontal eye fields. These connectivity peaks at anterior portions of the visual retinotopic maps suggest a broader linkage between areas at the boundary of classically-defined visual areas.

## Discussion

Integrating sensory and mnemonic information is a core problem faced by the brain(Rademaker et al., 2019; Favila et al., 2020; Groen et al., 2021; Rust and Palmer, 2021; Steel et al., 2024b). In the context of scenes, prior work has shown that co-localization of perceptual and memory-related brain areas facilitates their interaction: immediately anterior to the lateral and ventral scene perception areas exists a memory-responsive region that represents visuospatial context currently out of view(Silson et al., 2016b, 2019a; Steel et al., 2021, 2023, 2025; Srokova et al., 2022). Here, we show that the proposed fourth scene perception area in superior parietal cortex(Kennedy et al., 2024; Yoon et al., 2025) (SPA) also exhibits this motif. Using individualized fMRI stimuli and analyses, we show that SPA sits at the posterior edge of a large swath of cortex that responds selectively during recall of scenes (SPMA). Resting-state functional connectivity confirmed that SPA and SPMA differentially affiliate with the other scene perception and place memory areas, supporting roles in visual analysis versus representing visuospatial context, respectively. Finally, SPA sits at the intersection of two connectivity pathways: a medial pathway linking retrosplenial cortex and dorsal precuneus, and a lateral pathway linking caudal inferior parietal lobe to superior parietal and prefrontal cortex. These data suggest that SPA and SPMA may be uniquely positioned to coordinate mnemonic and perceptual reference frames to guide attention processes during visually-guided navigation.

Two studies have recently proposed adding a fourth region (SPA) to the set of brain areas considered important for scene perception (OPA, PPA, and MPA) (Kennedy et al., 2024; Yoon et al., 2025). Our data generally support the reported location and consistency of SPA. We observed SPA in the vast majority of our participants across two independent datasets, which used static (Dataset 1) and dynamic (Dataset 2) stimuli. In both datasets, this area was located at the dorsal extent of the parieto-occipital sulcus, separate from OPA. At the group level, SPA fell adjacent to the retinotopic map IPS1 in both datasets, and extended anteriorly into IPS2 in the group of participants who viewed dynamic stimuli. This anatomical location and preference for dynamic versus static stimuli agree with prior reports; however, our two datasets were collected independently, so we interpret this difference across datasets with caution. Notably, prior descriptions of PIGS located this area adjacent to the retinotopic IPS3 and IPS4(Kennedy et al., 2024), which is anterior and dorsal to our group analysis and more consistent with our memory-evoked activity. This discrepancy may be due to the more precise individualized retinotopic map definition in past work compared to our group atlas approach. Thus, while prior work investigating the neural basis of scene perception has focused on three areas (Epstein and Baker, 2019; Dilks et al., 2022), future work focused on visually-guided navigation may also incorporate SPA.

Using mental imagery of scenes to isolate top-down processes, we found that SPA falls at the posterior boundary of a larger memory-responsive cluster extending through precuneus toward the marginal ramus of the cingulate. We could localize this memory-responsive region, SPMA, in the majority of participants across two studies despite different tasks and experimental parameters, which speaks to its robustness. Like the other memory responsive regions (Steel et al., 2021, 2025), SPMA was anatomically distinct from its perceptual counterpart, with limited overlap between these functionally-defined areas. Interestingly, when viewing dynamic stimuli, SPMA retained its scene selectivity, suggesting that the preference for scenes in memory extends into perception in these areas at a sub-threshold level. Although this pattern may be driven by the small overlapping area between these functional regions of interest, further work should continue to resolve the functional dissociation of the perception and memory areas. The ability to localize SPA and SPMA in individual participants suggests that they are meaningful features of brain organization(Petersen et al., 2024) and opens up the possibility of studying these functional areas’ specific roles in coordinating scene perception and visuospatial memory.

The distinction between perception and memory is largely consistent on the lateral, ventral, and superior parietal surfaces. The medial surface responds strongly to mnemonic and spatial context, including familiarity with scenes and spatial knowledge (Bar and Aminoff, 2003; Sugiura et al., 2005; Epstein et al., 2007a, 2007b; Park and Chun, 2009; Robertson et al., 2016; Persichetti and Dilks, 2018, 2019; Berens et al., 2021). One possibility is that this surface constitutes a unified perception-memory architecture rather than a topographically distinct pairing. We did not observe a consistent anterior shift from perception to memory on the medial surface, consistent with other work that has also failed to observe this effect (Steel et al., 2025), and our prior work found similar responses to increasing spatial knowledge during perception and imagery in MPA and MPMA (Steel et al., 2023). A second possibility is a more complex distinction not well captured by the simple anterior-shift heuristic. For example, other work has observed a perception-memory distinction on the medial surface, including distinct peaks in activity across the boundary of the parieto-occipital cortex (Silson et al., 2019a) and intermixed positive and negative retinotopic coding in this area (Scrivener and Silson, 2025). Resolving this will likely require targeted investigation of medial parietal cortex, including tasks designed specifically to engage this region and high-resolution fMRI to disambiguate the anterior and posterior banks of the parieto-occipital sulcus.

What navigation processes might the scene perception and memory areas in superior parietal cortex serve? Although SPA has only recently been added to the “scene perception areas”, prior work has implicated superior parietal cortex in navigationally-relevant processes. Patients with superior parietal lesions exhibit “egocentric disorientation,” characterized by inability to locate objects and navigational affordances relative to themselves(Aguirre and D’Esposito, 1999). In addition, fMRI studies have noted superior parietal activation during egocentric navigationally-relevant tasks (Sulpizio et al., 2023), including integrating information across changes viewpoint (Van Assche et al., 2014), evaluating paths through a scene (Persichetti and Dilks, 2018), judging relative directions between viewpoints from memory (Marchette et al., 2014; Huffman and Ekstrom, 2019), and actively navigating (Suzuki et al., 2021). One study that employed a similar memory localizer task to our Edinburgh dataset found that superior parietal cortex appeared to represent integrated local viewpoints (Han and Epstein, 2026). In addition, two recent studies implementing an encoding model approach to fMRI data analysis provide additional, fine-grained insight into superior parietal cortex’s role in navigation (Lu et al., 2025; Zhang et al., 2025). One elegant study employing a navigation task in identical but visually-distinct environments observed precise and unique head direction coding in superior parietal cortex and in retrosplenial complex/MPA (Lu et al., 2025). A similar study with a large navigable environment and an elaborate set of navigation-relevant analytical regressors also observed that head-direction coding in superior parietal cortex, along with a mixture of motion energy, gaze semantics, and future path (Zhang et al., 2025). Our work adds to this evidence, showing that simply visually recalling navigationally-relevant information from an egocentric perspective is sufficient to activate superior parietal cortex.

SPA sits at a junction of four distinct connectivity branches: two medial branches between the parieto-occiptal cortex and dorsal precuneus, and two lateral branches between lateral occipital and parietal cortices. These branches were bordered on by two parallel streams of connectivity to SPMA, with each the SPMA stream’s endpoint falling anterior to SPA’s. This finding reinforces two key points. First, these results show that perception and memory processes are intimately paired throughout cortex, and superior parietal cortex appears to be an area in which these processes come together. Second, they further support the expanded role of superior parietal cortex in navigation function. Importantly, these branches span regions implicated in navigation-relevant processes, including processes related to allocentric map-like navigation(Marchette et al., 2014; Persichetti and Dilks, 2018; Lu et al., 2025; Naveilhan et al., 2025), perspective taking(Zhang et al., 2025), and ego-centric scene perception(Pitcher et al., 2011; Persichetti and Dilks, 2016, 2019). Being situated at the intersection of these streams puts superior parietal cortex, home to SPA and SPMA, in a unique position as an integrative hub, where our current percept is connected with our internal representation of location and current goal to guide our attention and inform our movement.

In summary, successful navigation requires both extracting visual information from the current scene and integrating that input with stored knowledge of the broader environment. A path is only viable if you know where it leads, and a landmark is only useful if you know what it signifies. Each scene perception area extracts a spatially-biased slice of the current visual scene (Silson et al., 2015, 2016a). However, that slice is only meaningful in light of context that extends beyond what is currently visible.

The architecture of scene processing cortex is organized to meet this demand: the scene perception areas are anatomically coupled with memory-responsive partners. This organization is particularly prominent on the lateral, ventral, and superior parietal surfaces, which clearly exhibit this posterior-anterior perception-memory motif. While the shift in activity is less prominent on the medial surface, memory recall evokes activity in a larger swath of cortex in this region, suggesting that MPA may be a visually-responsive subarea within a larger territory involved in memory-related processing. Crucially, each memory area connects to holistic representations of space in the hippocampus and our goals in prefrontal cortex. This may allow contextual information from the memory areas to befed back to shape ongoing perceptual processing (Angeli et al., 2024; Steel et al., 2024b, 2024a), such that what OPA “sees” as a viable path, or what PPA recognizes as a meaningful landmark, is jointly determined by current visual input and stored knowledge of the environment. In this sense, “visually-guided navigation” is not the function of a particular brain area — it emerges from the connection between each scene area’s visual computations and the context that gives them meaning.

## Acknowledgments

This project was funded by a Biotechnology and Biological Sciences Research Council award (UKRI2952) to ES, and by Neukom Institute for Computational Sciences to AS. AS also received funding from the Brain and Behavioral Research Foundation.

## Data availability

Data are available via OSF (Experiment 1 (Edinburgh): https://osf.io/ypk6c/; Experiment 2 (Dartmouth): https://osf.io/v4738/).

## Code availability

Analysis code is available via github (https://github.com/the-steel-lab/).

## Conflict of interest statement

The authors declare no conflict of interest.

## Author contributions

AS and ES conceived the study and designed the experiment. AS, LB, and CS processed the data. AS, NT, and DP analyzed the data. AS wrote the first draft. All authors edited and approved the final paper.

## Notes

### Competing Interest Statement

The authors have declared no competing interest.

### Summary of Updates

Simplified our ROI naming convention. We now refer to the areas as SPA (Superior place area) and SPMA (Superior place memory area), which better aligns with the larger scene perception literature. Added a discussion of the lack of topographic shift between MPA/MPMA. This included tempering our overall claim of ubiquitous shifting in the abstract and discussion. Added an analysis of the memory areas activity during perception. This revealed significant selectivity during perception of dynamic visual scenes. This complements the observation of anterior shifted memory activity. Added an analysis of connectivity from SPMA. This revealed an intriguing parallel between the streams of connectivity from SPA and SPMA. Report effect sizes and confirmed all statistical analyses.

